# Item recognition and lure discrimination in younger and older adults are supported by alpha/beta desynchronization

**DOI:** 10.1101/2020.04.22.055764

**Authors:** Anna E. Karlsson, Claudia C. Wehrspaun, Myriam C. Sander

**Affiliations:** Center for Lifespan Psychology, Max Planck Institute for Human Development, Berlin, Germany; Department of Psychology, Humboldt Universität zu Berlin, Germany

**Keywords:** aging, alpha/beta oscillations, EEG, episodic memory, lure discrimination

## Abstract

Our episodic memories vary in their specificity, ranging from a mere sense of familiarity to detailed recollection of the initial experience. Recent work suggests that alpha/beta desynchronization promotes information flow through the cortex, tracking the richness in detail of recovered memory representations. At the same time, as we age, memories become less vivid and detailed, which may be reflected in age-related reductions in alpha/beta desynchronization during retrieval. To understand age differences in the specificity of episodic memories, we investigated differences in alpha/beta desynchronization between younger (18–26 years, *n* = 31) and older (65–76 years, *n* = 27) adults during item recognition and lure discrimination.

Alpha/beta desynchronization increased linearly with the demand for memory specificity, i.e., the requirement to retrieve details for an accurate response, across retrieval situations (correct rejections < item recognition < lure discrimination). Stronger alpha/beta desynchronization was related to memory success, as indicated by reliable activation differences between correct and incorrect memory responses. In line with the assumption of a loss of mnemonic detail in older age, older adults had more difficulties than younger adults to discriminate lures from targets. Importantly, they also showed a reduced modulation of alpha/beta desynchronization across retrieval demands. Together, these results extend previous findings by demonstrating that alpha/beta desynchronization dissociates between item recognition and the retrieval of highly detailed memories as required in lure discrimination, and that age-related impairments in episodic retrieval are accompanied by attenuated modulations in the alpha/beta band. Thus, we provide novel findings suggesting that alpha/beta desynchronization tracks mnemonic specificity and that changes in these oscillatory mechanisms may underlie age-related declines in episodic memory.

## 1. Introduction

Episodic memory, the ability to remember events from the past, allows us to mentally travel back in time to re-visit places and people we have encountered (Tulving, 1985). It is crucial for human cognition and behavior, since it allows us to learn from experience and to detect novel elements in our surroundings. Memory retrieval can range from a mere sense of familiarity, that is, when we fail to recover specifying details but experience the event as familiar, to the recollection of highly detailed memories accompanied by the retrieval of specifying information (Jacoby, 1998; Mandler, 1980; for review, see Yonelinas, 2002). With advancing age, the ability to retrieve episodic memories declines (Naveh-Benjamin et al., 2003; Nyberg et al., 2012) and older adults’ memories seem to be less vivid and detailed (Folville et al., 2020; Greene & Naveh-Benjamin, 2020) than younger adults’. Higher levels of memory errors in older adults indicate the importance of retrieving memories with a high level of detail (Bowman et al., 2019; Fandakova et al., 2018; Koutstaal & Schacter, 1997). Nevertheless, the neural mechanisms underlying the loss of memory specificity with advancing age are still poorly understood.

A growing body of research has established the involvement of neural oscillations in memory processes (Düzel et al., 2010; Hanslmayr et al., 2016; Herweg et al., 2020; Nyhus & Curran, 2010). Brain oscillations provide a temporal window for the synchronous inhibitory or excitatory activity of neural cell assemblies. Such synchronous activity shapes synaptic weights within these assemblies (Markram et al., 1997) and thereby serves information processing related to memory formation and retrieval. Recent work has demonstrated that successful memory encoding and retrieval are associated with decreases in power in the alpha/beta frequency range (8–20 Hz; Griffiths et al., 2019; Waldhauser et al., 2016; for review, see Hanslmayr et al., 2012), reflecting the desynchronization of neural activity (Buzsáki et al., 2012). According to recent theoretical and empirical work, alpha/beta desynchronization is critical for efficient information processing (Hanslmayr et al., 2012; Parish et al., 2018) serving as a gating mechanism allowing for the flow of information through relevant neural assemblies. In support of this notion, alpha/beta desynchronization has been shown to reflect the level of processing at encoding (Hanslmayr et al., 2009; for review, see Hanslmayr & Staudigl, 2014) and seems to track the reactivation of mnemonic content at retrieval. Specifically, alpha/beta desynchronization was linearly related to the amount of information retrieved (Khader & Rösler, 2011; Waldhauser et al., 2016) and to the fidelity of information represented in cortex (Griffiths et al., 2019). Importantly, alpha/beta desynchronization does not seem to represent mnemonic information per se, but initiates neural conditions beneficial for cortical reinstatement, presumably by promoting information flow to the neural substrates conveying information about mnemonic content. Accordingly, two recent studies have reported that alpha/beta desynchronization increased systematically across conditions characterized by different levels of retrieval demands. In these studies, alpha/beta desynchronization showed a linear increase across item recognition and associative memory (Martín-Buro et al., 2020; Staresina et al., 2016), suggesting that the level of desynchronization might vary depending on the level of mnemonic detail needed for a correct response. In turn, the level of alpha/beta desynchronization may be considered as an index of the specificity of a recovered memory trace. However, the extent to which alpha/beta desynchronization is modulated by the amount of detail ingrained in a memory trace is still unclear, advocating the investigation of situations in which the retrieval of highly detailed, specific memories is needed, e.g., when the correct response depends on the detection of subtle changes in a test item.

The neural mechanisms supporting the ability to discriminate a similar lure from a previously encountered old item have typically been investigated in studies that extended the traditional old/new recognition memory paradigm by including similar lures in the test list, targeting the ability to remember details (Stark et al., 2019). These studies have provided evidence for the necessity to lay down orthogonalized memories in order to reduce interference, a process called pattern separation that relies primarily on the hippocampus (for reviews, see Stark et al., 2019; Yassa & Stark, 2011). However, other brain regions have also been shown to be important for the detection of similar lures. Specifically, regions throughout the ventral visual stream as well as earlier visual regions have been shown to dissociate between targets and perceptually and/or semantically similar lures (Bowman et al., 2019; Chouinard et al., 2008; Kim et al., 2009; Pidgeon & Morcom, 2016). Given the known role of these regions in the reactivation of mnemonic content (e.g., St-Laurent et al., 2015; Wing et al., 2015), successful target–lure discrimination presumably also relies on the recovery of fine-grained memory representations. As elaborated above, the level of alpha/beta desynchronization during memory retrieval may relate to the detailedness of episodic memories, in particular, and may reflect underlying differences in memory specificity. Thus, we predicted that alpha/beta desynchronization contributes not only to successful item recognition, as indexed by power differences between new and old information, but also supports the discrimination between highly similar mnemonic information underlying successful lure discrimination.

While the discrimination between highly similar mnemonic information is usually a difficult, yet feasible task for younger adults, older adults often show an impairment in tasks that require the retrieval of detailed information. Age differences are rarely observed when simple item recognition is sufficient, but older adults show difficulties when the retrieval of episodic detail is paramount to the task at hand, such as context or source information (e.g., Bastin et al., 2014; Luo & Craik, 2009; for review, see Old & Naveh-Benjamin, 2008). Similarly, the ability to discriminate similar lures from old items, which relies on the ability to recover memories with high specificity, is negatively affected by age (Stark et al., 2015; Stark & Stark, 2017; for review, see Stark et al., 2019) and is accompanied by an increased reliance on gist information and familiarity-based processes when making mnemonic judgments (Kensinger & Schacter, 1999; Koen & Yonelinas, 2014; Stark et al., 2015). The loss of mnemonic detail and reduced specificity of memory content in older adults is also reflected in age differences in the specificity of neural activation, so-called neural dedifferentiation (Bowman et al., 2019; Koen et al., 2019; Sommer et al., 2020; for review, see Koen & Rugg, 2019). Early functional magnetic resonance imaging (fMRI) studies demonstrated that while in younger adults, processing of category-specific information, such as faces or houses, is supported by category-selective regions (fusiform face area and parahippocampal place area respectively), older adults show a decrease in this neural selectivity (Park et al., 2004). Recent studies using multivariate pattern analysis support this initial finding and underline the relevance of neural specificity for memory performance (Kobelt et al. 2020; Koen et al., 2019; Zheng et al., 2018). Age differences in oscillatory mechanisms of memory formation (Werkle-Bergner et al., 2006) and retrieval may further contribute to the reduced memory specificity in older adults, but remarkably few studies have addressed this question as yet. In particular, given the evidence that the level of alpha/beta desynchronization during retrieval may track differences in memory specificity, a reduction in alpha/beta desynchronization in older adults during retrieval may be related to the observed memory decline. Among the very few studies investigating age differences in oscillatory underpinnings of episodic memory, a recent study showed that alpha/beta desynchronization during learning discriminates between subsequently remembered and subsequently forgotten item–picture associations in both younger and older adults (Sander et al., 2020; see also Strunk & Duarte, 2019). Importantly, the size of the subsequent memory effect depended on the structural integrity of the inferior frontal gyrus, a region that is known to be relevant for deep elaboration of information (Hanslmayr et al., 2011; Kim, 2011), and whose integrity was reduced in the group of older adults. Thus, the study by Sander and colleagues (2020) supports the notion that episodic memories differ between younger and older adults as a result of age differences during encoding (Craik, 2020). Regarding retrieval, alpha/beta desynchronization has been shown to distinguish between correct rejections of novel foils and correct pair memory judgments in both younger and older adults (Strunk et al., 2017). Together, these studies suggest that alpha/beta desynchronization, as a signature of the specificity of memories laid down at encoding and recovered at retrieval, represents a promising candidate for understanding age differences in episodic memory.

To summarize, we put forward the idea that the specificity of episodic memories determines the level of alpha/beta desynchronization during retrieval. Based on the observations that desynchronization in the alpha/beta frequency range is related to the level of processing at encoding (Hanslmayr & Staudigl, 2014) and to the quality of the recovered memory trace at retrieval (Griffiths et al., 2019), we hypothesized that also alpha/beta desynchronization at retrieval varies with retrieval demand, i.e. as a function of the level of mnemonic detail required to make an accurate memory judgment. The distinction between previously learned information and newly encountered information should be reflected in stronger alpha/beta desynchronization for old responses compared to new responses (see Martín-Buro et al., 2020). An even more fine-grained mnemonic representation is necessary when new information is highly similar to the learned information. Therefore, correct “similar” responses in a recognition memory task should be accompanied by even stronger alpha/beta desynchronization than correct “old” responses. Thus, we predicted a linear relationship between alpha/beta desynchronization at retrieval and the level of mnemonic detail required to make an accurate memory judgment (New < Old < Lure). In accordance with evidence for an age-related reduction in episodic detail of retrieved memories, we predicted age differences between younger and older adults when the discrimination between highly similar information is required, i.e., during lure discrimination. Furthermore, we hypothesized that older adults’ reduced memory performance would be accompanied by smaller modulations in alpha/beta desynchronization across experimental situations with varying retrieval demands. To this end, younger and older adults were measured with electroencephalography (EEG) while performing an episodic memory task. After studying pictures of everyday items, participants performed a surprise recognition memory test, where they were required to discriminate between old items, new items, and perceptually similar lures (see Fig. 1). This design allowed us to contrast neural activity between conditions with varying retrieval demands requiring different levels of mnemonic detail (correctly rejecting a new item, correctly remembering an old item, and correctly endorsing a lure as similar) in an age-comparative setting.

**Figure 1.**
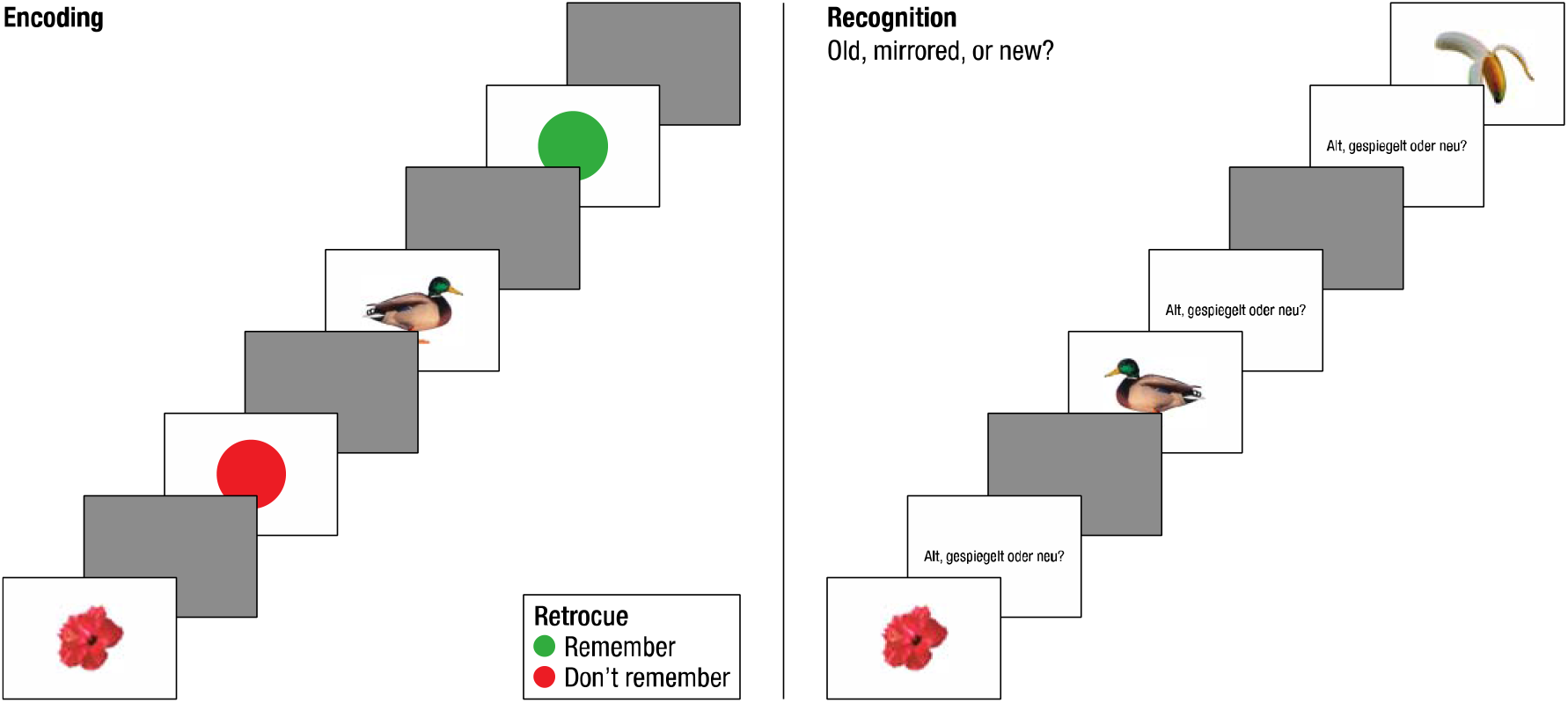
Experimental paradigm. During the encoding phase participants viewed different objects. Participants were instructed to either remember the object if it was followed by a green retro cue, or to not remember the object, if followed by a red retro cue. Objects were presented on flickering or white backgrounds. Note that neither the flicker nor the retro cue manipulations are relevant to the current analysis. After a 10-minute break, participants were asked to take a surprise memory recognition test. In this test, old objects either identical to the presentation during encoding or as their own mirror image (20% of the old objects) were presented together with completely new objects. Participants had to indicate whether the presented object was old, mirrored, or new.

## 2. Methods

### 2.1. Participants

Younger (*n* = 41) and older (*n* = 42) right-handed adults were recruited via a database of the Max-Planck Institute for Human Development to participate in the study for a compensation of €12/hour. One older and three younger adults were excluded due to behavioral performance below chance level (see section 2.5) and one older adult was excluded based on the MMST (Mini-Mental-Status Test; Folstein et al., 1975; see below). In addition, after preprocessing of the EEG data, another 7 younger and 13 older adults were excluded due to too few trials across the trial bins (see section 2.6). The remaining sample consisted of 31 younger adults (*M*_age_ = 21.84, *SD*_age_ = 2.6, range 18–26 years, 16 females) and 27 older adults (*M*_age_ = 70.74, *SD*_age_ = 2.73, range 65–76 years, 17 females).

To further characterize the sample, the participants completed a demographic questionnaire and a series of test batteries. Older adults were assessed using the MMST measuring cognitive impairment. A MMST score of 27 or higher indicates normal cognition, whereas a MMST score of 19–23 indicates mild cognitive impairment (O’Bryant et al., 2008). One older adult did not pass the cut off (MMST score = 23) and was excluded from further analysis. Although the MMST is not sufficient to with certainty rule out mild cognitive impairment, it is noteworthy that the remaining sample scored relatively high above the cut off (range: 28-30), and thus most likely represents relatively healthy older adults. All participants completed the Digit Symbol substitution test (DST; Wechsler, 1955). Younger adults showed a significantly higher DST score (*M* = 74.67; *SD* = 10.76) than older adults (*M* = 51.41; *SD* = 10.66; W = 787, α = 0.05, *p* < .001). Working memory (WM) was tested using the WM digit-sorting task (Kray, 2000), and younger adults (*M* = 9.77; *SD* = 2.64) did not score significantly higher than older adults (*M* = 8.44; *SD* = 3.04; W = 515, α = 0.05, *p* = .13). Performance in the marker tasks was similar to other cognitive neuroscience studies previously run at our research center (e.g., Fandakova et al., 2014; Sander et al., 2011) and showed a typical pattern of age differences. The ethics committee of the Max Planck Institute for Human Development, Berlin, approved the study.

### 2.2. Stimuli

The experiment and recording of behavioral responses were programmed in MATLAB (version 2016b; The MathWorks Inc., Natick, MA, USA), using the Psychophysics Toolbox (Brainard, 1997). The stimulus pool consisted of 250 real world asymmetric objects (500 x 500 pixels) from a publicly available data base (Brady et al., 2008). They were presented on the center of the screen.

### 2.3. Experimental paradigm

The experiment was performed in a dimly lit room that was electromagnetically and acoustically shielded. Prior to the task, participants received instructions and completed 15 practice trials that could be repeated if necessary. In addition, each experimental session started with written instructions on the screen and the participant initiated each session with a button press. Prior to starting the memory task, participants engaged in five minutes of wakeful rest. During encoding (Fig. 1), randomly drawn objects were presented against (*i*) a white background (50 stimuli), (*ii*) a background that flickered between white and black at a frequency of 36 Hz (50 stimuli), or (*iii*) a background that flickered between white and black at a frequency of 28.8 Hz (50 stimuli). Before the experiment, participants were informed of the flickering background and instructed to pay no attention to it. Effects of the flickering background at encoding are beyond the scope of this manuscript and will be ignored for the analysis of the recognition phase. However, to test for any effects of the flickering background on behavioral performance, we computed an ANOVA with flicker (36/28.8 Hz) and behavioral measure (lure discrimination/corrected recognition; see section 2.5 for the calculation of these measures) as within-subject factors, and age (older/younger) as between-subject factor. No significant main effect of (F(1,168) = 0.18, *p* = .675) or interactions with (*p* > .250) flicker were found. Therefore, the retrieval data were collapsed across flicker frequencies for the analysis.

Each encoding trial began with a black screen with a centered white fixation cross (henceforth referred to as ISI) with a jittered presentation time (1–1.5 s). Subsequently, the object was presented (2 s) followed by first another ISI (0.7 s) and second the presentation of a retro-cue (0.5 s), indicating either to remember (green dot), or not to remember (red dot) the previously shown item. The presentation times were chosen based on previous work that successfully used retro-cues (Mok et al., 2016). An ANOVA with cue (remember/don’t remember) and behavioral measure (lure discrimination/corrected recognition) as within-subject factors, and age (younger/older) as between-subject factor was used to assess whether cue condition had any effects on behavioral performance. No significant main effect of (F(1,168) = 2.99, *p* = .085) or interactions with (*p* > .320) cue were found and thus the data were collapsed across cue condition for the purpose of the current analysis.

After the encoding phase, participants completed a short sham recognition test aimed to prevent active rehearsal during the following 10-minute break. For ten randomly selected items (i.e., four new items and six old items that had been followed by a green retro cue), participants had to indicate whether the item was “old” (i.e., shown during the encoding phase), “mirrored” (i.e., the mirrored image was shown during the encoding phase), or “new” (i.e., the item had not been seen before). Following the sham recognition, subjects were randomly assigned to one of two different 10-minute break conditions, either engaging in wakeful rest (*n* = 30) while fixating a black screen with a centered white fixation cross or performing a distraction task (*n* = 28) during which they were instructed to imagine walking in a well-known public area in Berlin while imagining the scenery as vividly as possible. In line with previous evidence (Dewar et al., 2012), we expected higher performance after wakeful rest than after a distraction task. An ANOVA with behavioral measure (lure discrimination/corrected recognition) as within-subject factor, and rest (wakeful rest/distraction task) and age (younger/older) as between-subject factors, yielded no significant main effect of rest (F(1,54) = 0.82, *p* = .368) but a significant behavioral measure x rest x age interaction (F(1,54) = 5.74, *p* = .020, □^2^ = 0.10). However, no pairwise comparisons were significant (*p* > .083; see also Varma et al., 2017). We therefore collapsed across both groups in the current analysis.

Following the break, participants performed a recognition task including all remaining objects (n = 140) that had been shown during the encoding phase. These were randomly presented intermixed with pictures of new objects (*n* = 100). For 20% of the old objects (i.e., *n* = 28), the respective mirror image was shown and participants were asked to indicate by key press whether the object was (*i*) old, (*ii*) mirrored, or (*iii*) new (Fig. 1; Motley & Kirwan, 2012). Each trial started with a jittered ISI (1–1.5 s) followed by the presentation of an object (2 s). To prevent potential confounds of motor responses in the EEG signal, participants withheld their response until a subsequently presented screen provided the response alternatives and their mapping onto the cursor keys on the keyboard. Responses were made with the right hand and the next trial began when a response was given. Given that participants had to withhold their responses, the use of response times as indicators of cognitive dynamics is clearly limited. Accordingly, an ANOVA on median response times with retrieval condition as within-subject factor (old/new/lure) and age (younger/older) as between-subject factor yielded no significant effects (all *p*s > .067).

### 2.4. EEG recording and preprocessing

EEG data were continuously recorded with BrainVision Recorder (BrainVision Products GmbH, Gilching, Germany) using elastic caps with 61 Ag/Ag–Cl passive electrodes placed according to the 10-10 system. AFz was used as ground electrode and the right mastoid served as reference during recording. Additional electrodes were placed on the left mastoid for later re-referencing, and on the outer canthi (horizontal EOG) and below the left eye (vertical EOG) to monitor eye movements. Electrode preparation ensured impedances below 5 kΩ before recording. A band-pass filter of 0.1 to 250 Hz was applied and data were recorded with a sampling rate of 1000 Hz and a sampling interval of 1000 μS. During recording, EEG was monitored online, and channels with extremely noisy signal or excessive impedance were marked to be excluded from further analysis. In addition, heart rate was recorded via electrocardiogram (ECG) to remove possible cardiovascular artifacts from the EEG signal (see below).

The EEG data were preprocessed and analyzed with the FieldTrip toolbox (Oostenveld et al., 2011) and custom-written MATLAB code. Offline, EEG data were filtered (4^th^ filter order) with a pass-band of 1–125 Hz, and re-referenced to the linked mastoid channels. The ECG data were filtered with a band-stop filter (48–52 Hz) and appended with the preprocessed EEG data. Next, the data were segmented into 6-second epochs (−2000–4000 ms) and all trials were visually inspected. Trials with extreme artifacts were removed for the ICA only. Saccade-related transient spike potentials were identified using the COSTRAP algorithm and removed from the signal as independent components (Hassler et al., 2011). Blink, eye-movement, muscle, and heartbeat artifacts were detected using ICA (Bell & Sejnowski, 1995) and removed from the signal. Artifact-contaminated channels and trials (determined across all epochs) were automatically identified (1) using the FASTER algorithm (Nolan et al., 2010) and (2) by detecting outliers exceeding four standard deviations of the kurtosis of the distribution of power values in each epoch within low- (0.2–2 Hz) or high-frequency (30–100 Hz) bands respectively. Channels labeled as artifact-contaminated were interpolated using spherical splines (Perrin et al., 1989). Finally, a second visual inspection of each trial per participant was performed and any remaining artifact-contaminated trials were excluded from further analysis, resulting in an average of 15.3% of trials rejected. The preprocessed EEG data were subjected to time-frequency decomposition of each epoch (−2000–4000 ms, 50 ms sliding window) using a Morlet wavelet approach (wavelet width = 7) as implemented in FieldTrip, estimating spectral power from 1 to 20 Hz in steps of 1 Hz. Finally, data was segmented into the time-window of interest (0–2000 ms) and the power spectrum was log transformed on the single trial level (Smulders et al., 2018). Note that no baseline correction was applied to the data.

### 2.5. Behavioral analysis

To determine performance above chance level, a balanced accuracy score was calculated by taking the mean over the proportion of correct responses to new items, old items, and lures respectively, yielding a chance level of 33%.

Responses were categorized as *old hits* (responding “old” to an old item), *old misses* (“new”/”mirrored” to an old item), *lure hits* (“mirrored” to a lure), *lure misses* (“old” to a lure), *correct rejections* (CR; correctly rejecting a new item), and *false alarms* (FA; endorsing a new item as “old”). The different proportions of responses given to the three test item categories are presented in Fig. 2. To assess younger and older adults’ ability to recognize an old item while correcting for response bias, we computed corrected recognition scores by subtracting FA rates from old hit rates for each participant (Snodgrass & Corwin, 1988). Similarly, to evaluate participants’ abilities to discriminate mirrored lures from old items, we computed a lure discrimination index (LDI) by subtracting the lure miss rate from the lure hit rate (“Mirrored”|Lure – “Old”|Lure; Ngo et al., 2018; Toner et al., 2009).

**Figure 2.**
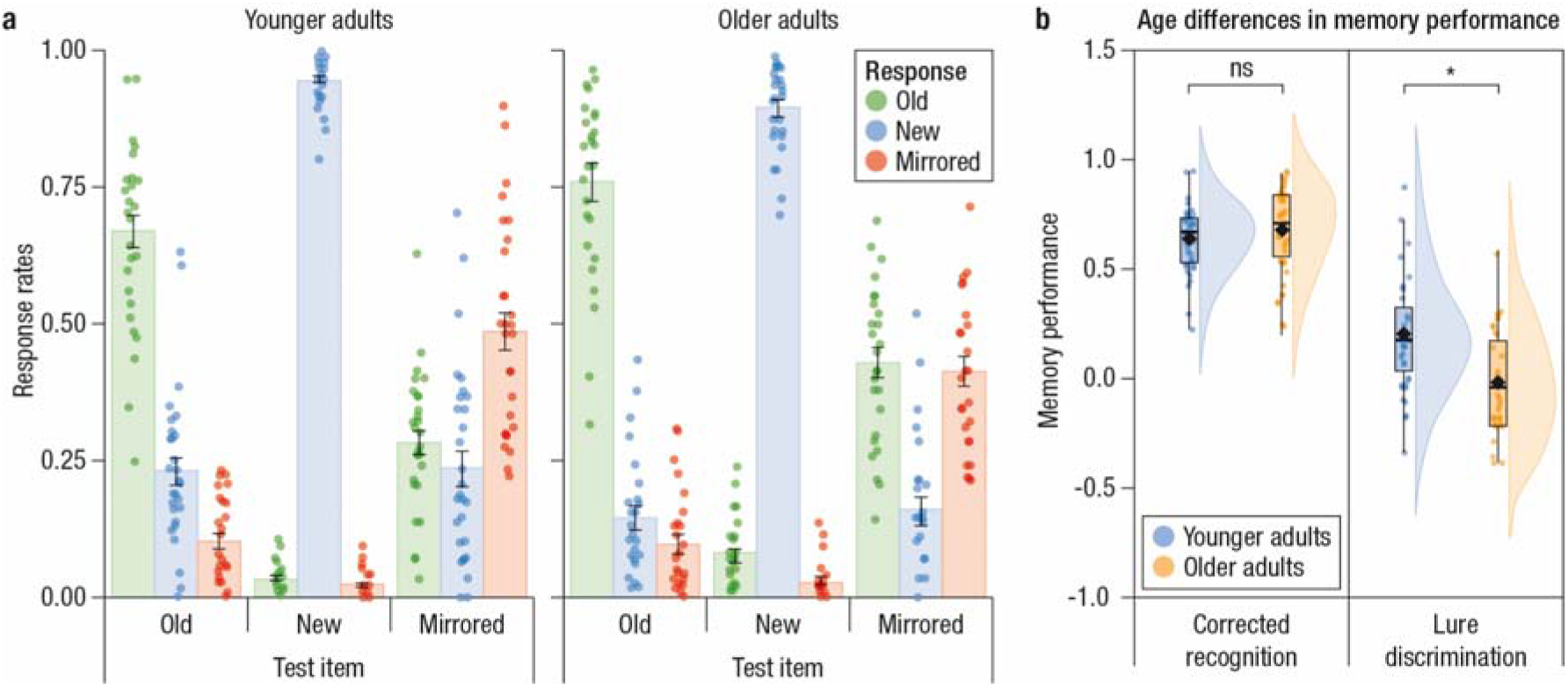
A: Response rates (y-axis) for the different responses, “Old” (green), “New” (purple), and “Mirrored” (red), across the different test item categories (x-axis) for younger (top) and older (bottom) adults. Bars represent the means with error bars showing standard deviations. Dots indicate individual data points. B: Illustration of the age difference evident for lure discrimination. Memory performance score (y-axis) for the two memory measures (corrected recognition, lure discrimination index; x-axis) across age groups (younger in blue, older in orange). Boxplots represent the interquartile range (first and third quantile) with dots representing individual participants. The rhombus indicates the mean, the horizontal bar indicates the median, and violin plots illustrate the sample density. The asterisk reflects significance below .05.

To investigate age differences in item recognition and lure discrimination, we performed a mixed effects ANOVA with age (younger/older) as between-subject, and measure (corrected recognition/LDI) as within-subject factors. Any significant interactions were followed up with pairwise comparisons. Bonferroni-corrected p-values are reported.

### 2.6. EEG group level analysis

To quantify the differences in power between conditions of different retrieval demands, trials associated with correctly rejecting a new item (CR), correctly recognizing an old item (old hits), and correctly discriminating a lure (lure hits), were binned and averaged within participants. Participants with less than a minimum of 6 trials per bin were excluded from the analysis (see section 2.1 for sample information), resulting in an average of 84.9 (range: 68–94) and 82 (range: 59–93) CR trials, 72.5 (range: 28–108) and 82 (range: 35– 109) old-hit trials, and 12.6 (range: 6–26) and 11.1 (range: 6–20) lure-hit trials for younger and older adults respectively.

The averaged time-frequency power values were subjected to a non-parametric, cluster-based, random permutation test (FieldTrip toolbox; Maris & Oostenveld, 2007), contrasting the time-frequency power spectrum between conditions with different retrieval demands (CR < old hits < lure hits) at each channel, frequency band (2–30 Hz), and time point (0–2 s). Initially, clusters were formed based on univariate two-sided, dependent samples regression coefficient *t*-statistics for each electrode–frequency–time point. The threshold for data points to be included in a cluster was set to *p* = .05 and the spatial constraint was set to a minimum of two neighboring channels. Next, the significance of the cluster-level statistic (i.e., the summed t-values) was assessed by comparison to a permutation null-distribution, which was obtained by randomly switching the condition labels and re-computing the *t*-test 1000 times. The final cluster *p*-value (i.e., the Monte Carlo significance probability) is the proportion of random partitions in which the cluster-level statistics were exceeded. The cluster-level significance threshold was set to *p*-values below .025 (two-sided significance threshold). This analysis yielded one significant negative cluster in the alpha/beta frequency range, comprising most channels from ~600–2000 ms (see section 3.2). This electrode–frequency–time cluster was used for the following subsequent analyses:

1. To explore potential age differences in the strength of the observed alpha/beta desynchronization effect (i.e., the slope), we computed individual linear regressions and averaged the individual regression coefficients within the electrode–frequency–time cluster that was identified across younger and older adults (see above). We tested for differences between the age groups using an independent samples *t*-test.
2. To examine the relevance of the level of alpha/beta desynchronization for memory performance, we contrasted neural activity associated with successful and unsuccessful responses for item recognition and lure discrimination. To this end, we extracted and averaged the individual log-transformed power values within the identified time–frequency–electrode cluster for each condition of interest (i.e., old hits, old misses, lure hits, and lure misses) and participant. Power values were entered in two ANOVAs with age (younger, older) as between-subject, and accuracy (Hit, Miss) as within-subject factors for old and lure items respectively. Significant interactions were followed-up with pairwise comparisons. Reported p-values are Bonferroni-corrected.

Note that while the electrode–frequency–time cluster was based on trials with correct responses only, this analysis contrasts correct and incorrect responses to assess the behavioral relevance of the effect. We would like to point out that whereas lure misses includes only “old” responses and thus allows us to dissociate lure discrimination from item recognition, old misses includes all incorrect responses (i.e., “new” and “mirrored”). Although “mirrored” and “new” responses to old items most likely involve different cognitive processes (i.e. familiarity and the absence of familiarity respectively; see Yonelinas, 2002), both responses are characterized by an absence of recollection. Thus, by categorizing “mirrored” and “new” responses as misses, the contrast aim to target pure recognition as compared to familiarity-based memory.

Some participants contributed fewer than 6 trials per condition and were excluded from this analysis, resulting in a sub-sample of 45 participants (23 younger adults, 22 older adults) with an average of 75.3 (range: 47–108) and 82.3 (range: 35– 105) old-hit trials, 33 (range: 6–62) and 27.4 (range: 6–76) old-miss trials, 11.7 (range: 6–20) and 10.6 (range: 6–17) lure-hit trials, and 8.9 (range: 6–17) and 12.7 (range: 7–20) lure-miss trials for younger and older adults respectively.

#### Data Accessibility

Data and custom Matlab code of the main analyses are available on the Open Science Framework repository (https://osf.io/dt4ax/).

## 3. Results

### 3.1. Older adults show reduced lure discrimination

To investigate age differences in item recognition and lure discrimination, a mixed ANOVA with age (younger, older) as between, and performance measure (corrected recognition, LDI) as within-subject factors was implemented. The results (see Fig. 2) demonstrated a marginally significant main effect of age (F(1;56) = 3.82; *p* = 0.056, *np2* = 0.1) and a main effect of measure (F(1;56) = 249.19; *p* < .001, *np2* = 0.82), reflecting lower performance in older (*M* = 0.33, *SD* = 0.41) than younger (*M* = 0.42, *SD* = 0.31) adults as well as better recognition memory (*M* = 0.66, *SD* = 0.18) than lure discrimination (*M* = 0.1, *SD* = 0.28). In addition, a significant age x measure interaction was found (F(1;56) = 13.81; *p* < .001, *np2* = 0.2), reflecting comparable recognition memory across age groups (*t*(51.8) = −0.9, *p* > .250) but reduced lure discrimination in older compared to younger adults (*t*(55.71) = 3.3, *p* = 0.003, d = 0.86).

### 3.2. Alpha/beta desynchronization varies as a function of retrieval demand

Time-frequency representations (Fig. 3A) of the retrieval-related neural activity averaged over posterior channels (TP7, CP5, CP3, CP1, CPz, CP2, CP4, CP6, TP8, P7, P5, P3, P1, Pz, P2, P4, P6, P8, PO7, PO3, POz, PO4, PO8, O1, Oz, O2) display a clear increase in the theta frequency range (~3–7 Hz) following stimulus onset. This early increase in power is followed by a strong power decrease in the alpha/beta frequency range (~8–20 Hz). A similar oscillatory pattern is evident across the three conditions with different retrieval demands. In order to test for differences in alpha/beta desynchronization between the conditions, we conducted linear regressions across levels of retrieval demand (correct rejections < item recognition < lure discrimination), collapsed across age groups. A negative cluster was identified, reflecting a reliable linear relationship between alpha/beta desynchronization and retrieval demand (*p* < .001; see Fig. 3B & D). This negative cluster showed a wide-spread topographical distribution with a maximum over posterior channels (see Fig. 3C), temporally encompassed most of the epoch (~0.6–2 s), and comprised a large frequency range within the alpha and beta frequency bands (~8–25 Hz). To assess age differences in the strength of the observed alpha/beta desynchronization effect (i.e., the slope), we compared younger and older adults’ regression coefficients within the electrode–frequency–time cluster. Importantly, older and younger adults differed with regard to the strength of the effect (*t*(54.51) = −2.35, *p* = 0.022, d = 0.62), reflected in a steeper slope in younger (*M* = −0.82, *SD* = 0.41) as compared to older (*M* = −0.56, *SD* = 0.42) adults (see Fig. 3E).

**Figure 3.**
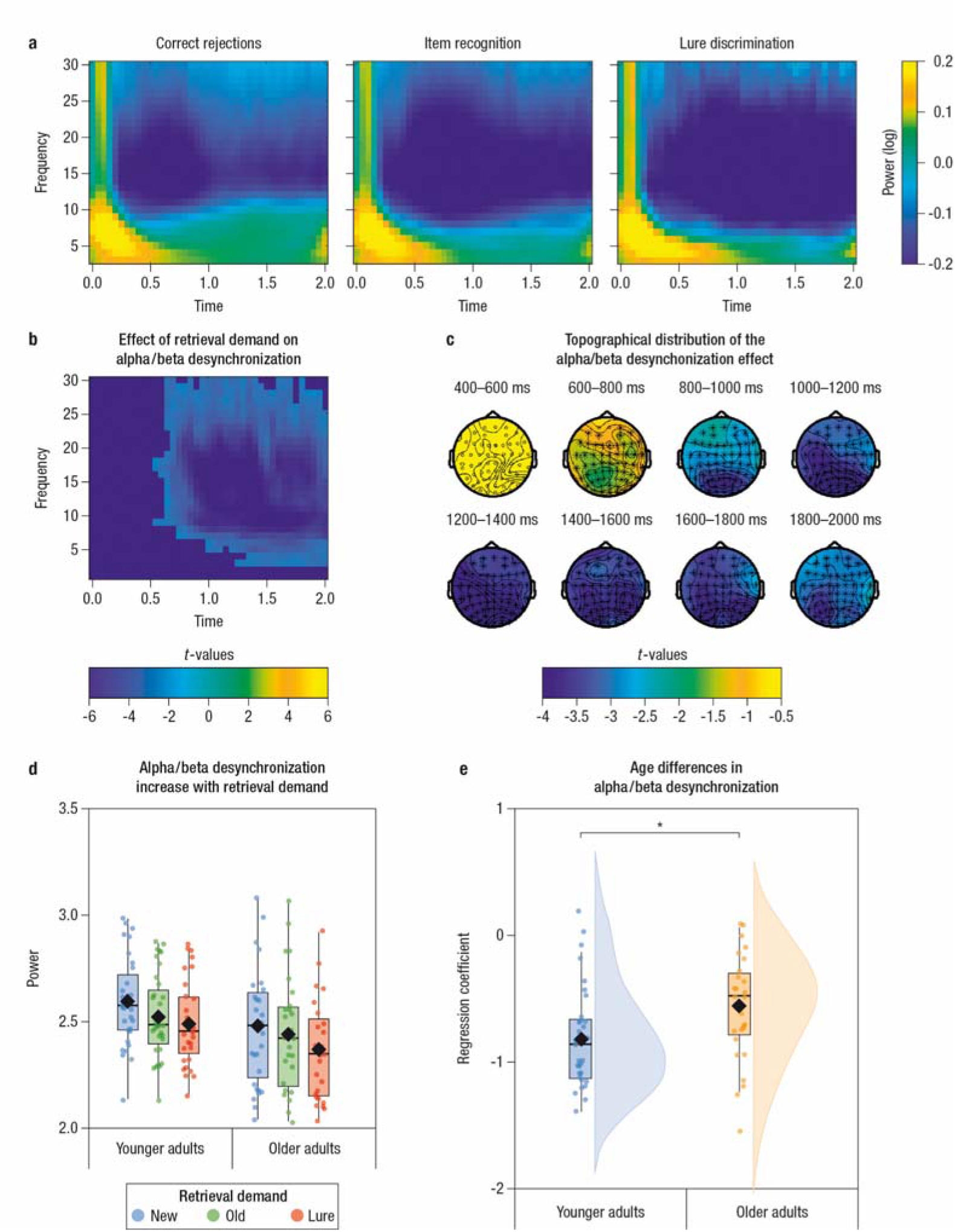
A: Time (x-axis) – frequency (y-axis) representations of log-transformed power for conditions of different retrieval demands (correct rejections, item recognition, lure discrimination) averaged over participants. Note that the data is plotted with baseline correction (absolute baseline, −0.5–0.2 s) for illustrative purposes only. For analysis, no baseline correction was applied. B: Results of the cluster-permutation analysis using dependent regression to test for differences in alpha/beta power across retrieval conditions. The highlighted negative time-frequency cluster indicates a reliable linear increase in alpha/beta desynchronization across retrieval demands. C: Topographical distribution of the effect from 400-2000 ms in steps of 200 ms. The effect is wide-spread across all channels but with a posterior maximum throughout the epoch. The asterisks highlight significant channels. D: For visualization purposes only, we display log-transformed power (y-axis) averaged within the negative cluster by age group (x-axis). This illustrates that the linear increase in alpha/beta desynchronization across retrieval demands is evident in both age groups. However, as displayed in (E): the slope of alpha/beta desynchronization (expressed as t-values of the individual linear regression averaged within the cluster) is significantly steeper in younger (blue) as compared to older (orange) adults. Boxplots represent the interquartile range (first and third quantile) with dots representing individual participants. The rhombus indicates the mean, the horizontal bar indicates the median, and violin plots illustrate the sample density. The asterisk reflects significance below .05.

### 3.3. Larger alpha/beta desynchronization predicts within-person memory performance

If the observed modulation of alpha/beta desynchronization by retrieval demand and age group contributes to memory performance, we would expect it to vary with regard to memory success. Therefore, we contrasted log-transformed power values within the observed cluster for successful versus unsuccessful item recognition as well as successful and unsuccessful lure discrimination. To this end, we conducted two separate ANOVAs, including age group (younger, older) as between-subject, and accuracy (Hits, Misses) as within-subject factors, for old and lure items respectively (see Fig. 4). For old items, a main effect of accuracy (F(1;43) = 6.723; *p* = 0.013, *np2* = 0.14) was found, indicating stronger alpha/beta desynchronization for hits (*M* = 2.48, *SD* = 0.24) as compared to misses (*M* = 2.5, *SD* = 0.25) regardless of age. However, a significant interaction of age and accuracy (F(1;43) = 10.32; *p* = 0.002, *np2* = 0.19) followed up with pairwise comparisons revealed that alpha/beta desynchronization was stronger for old item hits as compared to old item misses in younger adults (*t*(22) = −5.02, *p* < .001, d = 0.19), whereas no significant difference was evident for older adults (*t*(21) = 0.42, *p* > .250). For lure items, we found a main effect of accuracy (F(1;43) = 37.266; *p* < .001, *np2* = 0.46), reflecting stronger alpha/beta desynchronization for hits (*M* = 2.42, *SD* = 0.22) as compared to misses (*M* = 2.48, *SD* = 0.25) regardless of age. The interaction between age group and accuracy was not significant (F(1;43) = 2.29; *p* > 0.137).

**Figure 4.**
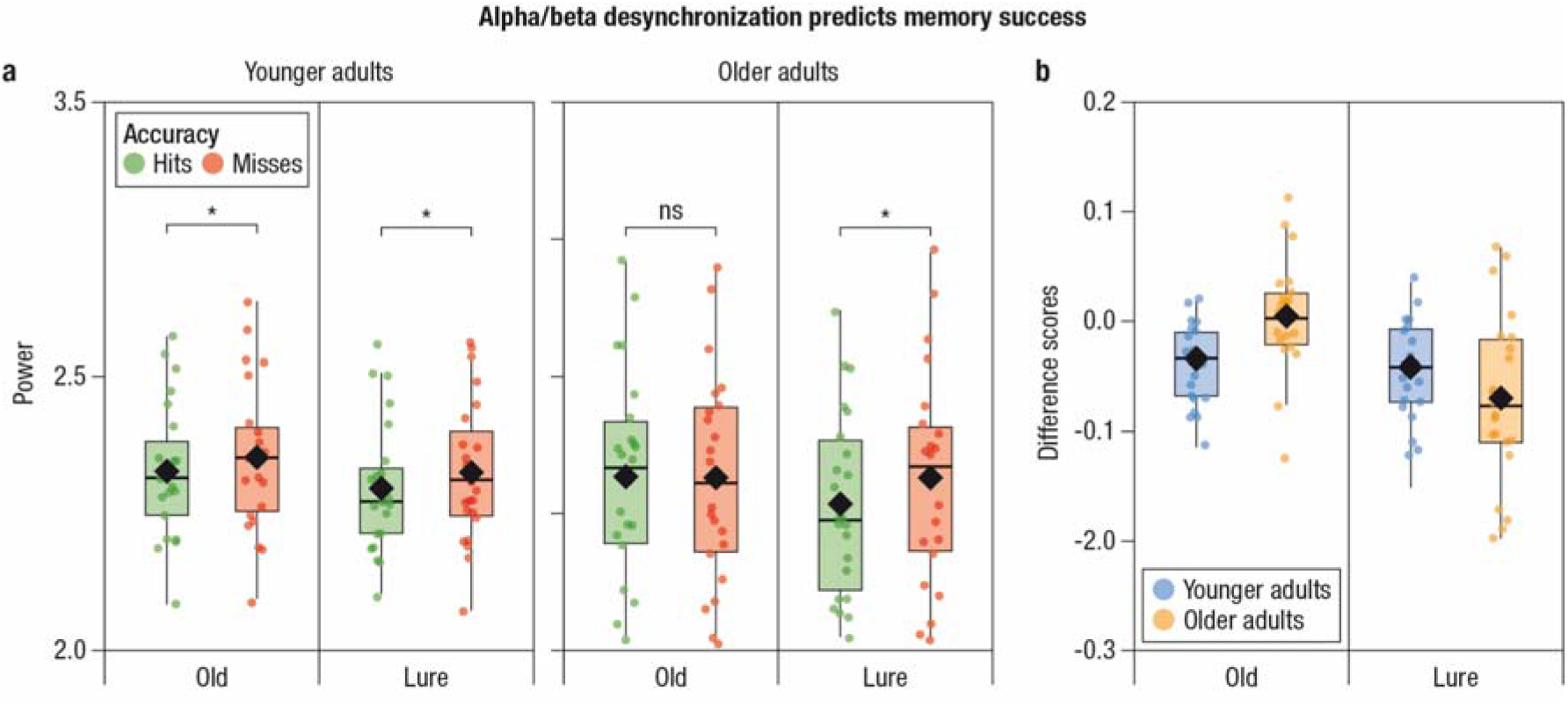
A: Log-transformed power (y-axis) averaged within the previously determined cluster (see Fig. 3B) by age group (left: younger adults, right: older adults) and accuracy (Hits: green, Misses: red). B: For illustrative purposes only, difference scores (Hits – Misses) as a function of retrieval demand (Old, Lure; x-axis) and age (younger: blue; older: orange), demonstrates the age differences in the modulation of alpha/beta desynchronization by retrieval success across retrieval conditions. Boxplots represent the interquartile range (first and third quantile) with dots representing individual participants. The rhombus indicates the mean and the horizontal bar indicates the median. Asterisks reflect significance below .05.

## 4. Discussion

Memory performance depends on the encoding and retrieval of detailed information. However, our memories are not all equally precise, but vary in their level of specificity, determining memory success and failure. Becoming older is accompanied by an increased loss of mnemonic specificity which may contribute to the commonly observed episodic memory decline (Nyberg et al., 2012; Wang et al., 2017). In light of recent work proposing that alpha/beta desynchronization relates to the quality or specificity of retrieved memories (Griffiths et al., 2019; Khader & Rösler, 2011; Martín-Buro et al., 2020), we investigated whether alpha/beta desynchronization was modulated by different retrieval demands in younger and older adults. Specifically, we asked whether the level of alpha/beta desynchronization is not only modulated by the need to detect old (compared to new) information as in successful item recognition, but also varies with the level of detail required for the discrimination between targets and lures, which provide highly similar information.

### 4.1. Alpha/beta desynchronization supports item recognition and lure discrimination

As hypothesized, alpha/beta desynchronization showed a linear relationship with retrieval demand, that is, the amount of detail required to make an accurate mnemonic judgment (New < Old < Lure; see Fig. 3.B). This finding supports the notion that the level of alpha/beta desynchronization reflects the level of mnemonic evidence available during retrieval and can serve as an index of memory specificity. Our findings parallel those of a recent study with a sample of young adults only by Martín-Buro and colleagues (2020). That study showed that alpha power decreases across retrieval conditions that differ in the amount of information needed for a correct response (correct rejections < item recognition < associative recall). Using source localization, the authors were able to link the observed alpha desynchronization to the posterior parietal cortex (PPC). Accordingly, they proposed that parietal alpha desynchronization may reflect the accumulation of mnemonic evidence governed by the PPC, serving as an ‘episodic buffer’ representing recollected information (Rugg & Vilberg, 2013). Although the effect reported here shows a widespread topography, it is seemingly maximal over posterior channels (see figure 3.C). Thus, our results extend these findings by demonstrating that alpha/beta desynchronization discriminates between item recognition and lure discrimination in a similar manner. Since target–lure discrimination depends on the detection of subtle differences, the retrieval of a highly detailed, specific memory representation is necessary for a correct memory judgement. We suggest that this higher memory specificity is supported by stronger alpha/beta desynchronization in lure conditions, indexing the larger amount of detail engrained in the recovered memory trace. Importantly, by contrasting trials associated with correct and incorrect memory responses we demonstrated reliably stronger desynchronization for correct as compared to incorrect retrieval. Thus, we expand previous research further by providing evidence that this gradual increase in alpha/beta desynchronization contributes to memory performance and predicts memory success both during item recognition and during discrimination of highly similar lures. In particular the difference in desynchronization between correct and incorrect lure responses provides further support for the view that alpha/beta desynchronization tracks the level of detail of retrieved memories (Hanslmayr et al., 2016), rather than the disinhibition of task relevant networks as posed by competing models of the functional role of alpha/beta oscillations (Jensen & Mazaheri, 2010; Klimesch et al., 2007). Given that incorrect trials were defined as “old” responses to lures, this contrast builds on the fine difference between the retrieval of the target memory without sufficient detail to detect the perceptual difference of the mirrored image, and the retrieval of a fine-grained memory representation. Thus, together with previous observations, our findings provide compelling evidence that alpha/beta desynchronization supports the gradual accumulation of mnemonic evidence and supports the successful recovery of highly detailed memories.

The observed effect of alpha/beta desynchronization during retrieval also parallels findings from earlier event-related potential (ERP) studies that aimed to dissociate memory processes associated with successful and unsuccessful lure discrimination. In particular, a left-parietal old/new effect (late positive potential: LPP) has been established as a neural correlate of successful recollection (see Rugg & Curran, 2007 for review). Some studies observed the LPP only for true memories (e.g., Curran, 2000; Poch et al.,2019), suggesting that lure false alarms are driven by familiarity. However, others have demonstrated that the LPP is observable during lure discrimination independent of memory success (e.g., Curran & Cleary, 2003; Morcom, 2015), suggesting that lure discrimination requires the recovery of mnemonic detail and evokes recollection. While we cannot clearly dissociate the relative contribution of familiarity and recollection in the present paradigm, the onset of the present effect falls within a similar time range (500–800 ms) as the well-established ERP marker of recollection and demonstrates a maximum over posterior channels, supporting our interpretation of alpha/beta desynchronization during retrieval as an index of the specificity of the retrieved content.

However, it is important to note that with the current design we cannot dissociate processes related to the recovery of mnemonic content, i.e. retrieval itself, and processes operating on retrieved content, such as cue specification and monitoring of retrieved content (for reviews on the involvement of the PFC in episodic memory see e.g. Simon & Spiers, 2003; Eichenbaum, 2017). Thus, it is possible that the effect observed here involves a mixture of these processes unfolding over time and jointly contributing to the accumulation of mnemonic evidence.

Previous work has highlighted the crucial role of the hippocampus for successful lure discrimination. The hippocampus is proposed to lay down non-overlapping memories during learning (cf. Stark et al., 2019, for review) and to prompt the recovery of the cortical memory traces at retrieval (Norman & O’Reilly, 2003; Rissman & Wagner, 2012). Accordingly, a recent intracranial EEG study (Griffiths et al., 2019) demonstrated that cortical alpha/beta desynchronization (in the anterior temporal lobe) was preceded by hippocampal gamma increases by a few hundred milliseconds during retrieval, suggesting a directionality in the flow of information from the hippocampus to the cortex. Furthermore, it is important to note that although the effect reported here were prominent in the alpha/beta band, activity in other frequency bands, especially in the theta frequency range (~3–7 Hz), have been shown to play a crucial role in memory processes. For instance, theta power increases have been associated with recollection (Gruber et al., 2008) and with hippocampal connectivity with cortical regions during retrieval (Herweg et al., 2016; for reviews, see Herweg et al., 2020; Nyhus & Curran, 2010; Staresina & Wimber, 2019). Thus, the recovery of detailed mnemonic content may rely on the interactive contribution of discrete oscillatory mechanisms, orchestrating the communication of information through functionally distinct parts of the episodic memory network. It seems that alpha/beta desynchronization supports the efficiency by which mnemonic information, as initiated by the hippocampus, is conveyed throughout the network and represented in ‘associative hubs’ such as the PPC (Rugg & King, 2017; Rugg & Vilberg, 2013), thereby ultimately supporting successful item recognition and lure discrimination.

### 4.2. Reduced alpha/beta desynchronization in older adults may indicate a loss in memory specificity

The strength of the observed alpha/beta desynchronization effect across retrieval demands (i.e., the slope) differed between age groups (see Fig. 3.E), with a smaller effect in older adults. This age difference in the modulation of alpha/beta power was accompanied by differences in lure discrimination abilities between younger and older adults (see Fig. 2), pointing to the potential contribution of age-related changes in alpha/beta desynchronization to episodic memory decline. Our findings are in line with converging evidence of detrimental effects of aging on the ability to retrieve episodic details (Bowman et al., 2019; Stark et al., 2019). We add to this evidence by showing that older adults’ difficulties to discriminate between highly overlapping mnemonic information may relate to less efficient desynchronization in the alpha/beta frequency bands, presumably reflecting less stable recovery of detailed memories. We further show that, in contrast to younger adults, alpha/beta desynchronization in older adults distinguished between correct and incorrect memories only when reliance on highly detailed memories, i.e., recollection, was required (lure items), but not when a correct response could rely on familiarity alone, i.e., in the case of old items (see Fig. 4). When older adults successfully identified a similar lure, alpha/beta desynchronization was stronger as compared to when they incorrectly endorsed a similar lure as old (and this effect was comparable across age groups). However, no differences in alpha/beta desynchronization were observed between correct and incorrect responses to old target items. Thus, we speculate that less efficient alpha/beta desynchronization in the aging brain indicates a reduction in information flow (Hanslmayr et al., 2012) which promotes impairments in the retrieval of detailed memories. As a consequence, older adults are more likely to rely on gist information and familiarity-based processing, making them more prone to incorrectly endorse a similar lure as an old item.

In the present study, we did not disentangle the degree to which age differences occurring at retrieval are a consequence of age differences during encoding. Recent findings suggest that age differences in episodic memory could also be attributed to encoding-related neural activity modulated by differences in the integrity of functionally relevant brain regions (Sander et al., 2020), and that memory quality (defined based on encoding success) influences retrieval-related processes differently in younger and older adults (Fandakova et al., 2018). Thus, it is possible that the age differences observed here are downstream effects of impoverished encoding rendering constructed memories less detailed. Investigations of the relation between alpha/beta desynchronization and the quality of represented mnemonic content across encoding and retrieval (see Griffiths et al., 2019 for an approach in this direction in younger adults) would be a valuable next step for future studies. While in the current study, we can only infer the level of mnemonic detail available to a person from the probability of correct responses to targets and lures, we propose that future studies could map within-person variations of single-trial alpha/beta power in younger and older adults directly to the representational properties of neural patterns of stimulus-related activity, for example, their distinctiveness (Koen & Rugg, 2019).

Finally, it is important to note that not only differences in retrieval, but also post-retrieval processes may have contributed to the observed age differences. The need to recruit monitoring processes increase with retrieval difficulty (e.g. Dobbins et al., 2002), and older adults show reductions in the recruitment of these processes, especially when evaluating information highly similar to previously learned material (e.g. Gutchess et al., 2007; Pidgeon & Morcom, 2014; Trelle et al., 2019). As in the present design, the lures are perceptually very similar to the target items, they potentially cause interference and require additional control processes to make a judgment. Thus, we cannot rule out the possibility that the age differences reported here result from the decline of a combination of processes that are necessary for successful lure discrimination, e.g. recovery of a highly detailed memory representation as well as monitoring of the retrieved content (e.g. Trelle et al., 2017; see Devitt & Schacter, 2016 for review).

To summarize, we showed that alpha/beta desynchronization increases across retrieval situations in which correct memory responses require different levels of detail. Age-related impairments in the retrieval of highly detailed memories were accompanied by reduced modulations of alpha/beta desynchronization. We further demonstrated that the level of alpha/beta desynchronization was predictive of memory success: In young adults, trials with correct and incorrect memory judgements differed reliably for both old targets and lures. However, in older adults, these effects of memory accuracy were only observed for lures, thus, when memory success depends on the retrieval of precise details. Taken together, our results suggest that alpha/beta desynchronization tracks the specificity of retrieved memories. Importantly, our findings also provide support for the notion that age differences in oscillatory mechanisms of retrieval contribute to age-related declines in episodic memory.

## Acknowledgements

This research was conducted within the “Lifespan Age Differences in Memory Representations” (LIME) project at the Center for Lifespan Psychology, Max Planck Institute for Human Development, Berlin, Germany. MCS was supported by the Minerva program of the Max Planck Society.

AEK is a doctoral fellow of the International Max Planck Research School on the Life Course (LIFE). We thank Gabriele Faust (LIME) and all student assistants and interns who helped with participant recruitment and data collection, project members for helpful feedback on the analysis, and all study participants for their time. We thank Julia Delius for editorial assistance.

